# Excessive zinc reduces histone acetylation in mouse cardiomyocytes by downregulating acetyltransferases

**DOI:** 10.1101/2025.03.09.642293

**Authors:** Shu Xu, Yuzhuang Hu, Chao Tang, Weize Xu

## Abstract

Zinc plays a critical role in cellular functions, but its excess may lead to detrimental effects, including cardiac abnormalities. The impact of zinc overload on cardiomyocytes is investigated in this study. For the first time, we report the regulatory relationship between zinc and histone acetylation. Excessive zinc downregulates the transcription of *Bmp4* in HL-1 cell line and primary cardiomyocytes. This downregulation is linked to reduced histone acetylation (H3K9ac), mediated by the suppression of histone acetyltransferases (HATs), rather than changes in histone deacetylases (HDACs). When zinc is introduced directly into the nucleus, *Bmp4* expression is upregulated, suggesting that the reduction of histone acetylation by zinc is in an indirect way. This study underscores the importance of zinc homeostasis in maintaining cardiac health and provides insights into the molecular basis of zinc-induced cardiac dysfunction.

## Introduction

Zinc is the second most abundant trace element in the human body after iron [1]. As early as 1961, zinc had already been identified as an essential trace element for humans [2]. A significant number of proteins in mammals contain zinc, with approximately 10% of human proteins being zinc proteins [3]. The homeostasis of intracellular zinc is crucial for maintaining the balance between health and disease. Long-term exposure to environmental zinc, such as through air, water, soil, and food, is detrimental to the digestive, respiratory, and nervous systems [4]. Excessive zinc disrupts energy production in the mitochondria of nerve cells, specifically manifested by increased production of reactive oxygen species (ROS), loss of mitochondrial membrane potential, and reduced cellular ATP levels [5].

Clinical reports also suggest that excessive zinc may lead to abnormal cardiac responses, including tachycardia, arrhythmia, and hypotension [6-9]. However, research on the corresponding pathogenic mechanisms remains relatively limited. Therefore, studying the molecular-level changes in cardiac cells under conditions of zinc overload holds significant medical importance.

This study utilized mouse cardiomyocytes as a model to investigate the impact of zinc overload on the transcriptional regulation of key cardiac factors, thereby uncovering the link between zinc and myocardial epigenetic. For the first time, our study discovered that zinc can reduce acetylation levels in cardiomyocytes by regulating the transcription of acetyltransferases, subsequently affecting the transcriptional expression of a wide range of downstream genes. This study provides a molecular-level understanding of the symptoms caused by the imbalance of myocardial trace element homeostasis in clinical settings.

## Materials and Methods

### Cell Lines

Mouse cardiac muscle cell line HL-1 was obtained from Sigma-Aldrich (SCC065). Cells were cultured in a Claycomb medium (51800C, Sigma-Aldrich) supplemented with 10% fetal bovine serum (FBS) (Life Technologies, Inc., Grand Island, NY), 1% penicillin/streptomycin, Norepinephrine 10 mM (A0937, Sigma-Aldrich), and L-Glutamine 2 mM (A7506, Sigma-Aldrich). AC16 human cardiomyocyte cell lines were purchased from Sigma-Aldrich (SCC109). Cells were cultured in high glucose DMEM (Life Technologies, Inc., Grand Island, NY) containing 10% fetal bovine serum (FBS) (FBS, Life Technologies, Inc., Grand Island, NY), 1% antibiotics penicillin–streptomycin. All the cells were maintained at 37 °C with 5% CO_2_.

### Cell viability assay

Cell viability was assayed by Cell Counting Kit-8 (HY-K0301, Med Chem Express) according to the manufacturer’s protocols. Briefly, HL-1 and AC16 cells were seeded into 96-well plates and cultured for 12 hours. After treatment for 24 hours, 10 μL of CCK-8 solution was added to each well and incubated for 2 h at 37 °C. The absorbance was then recorded at 450 nm using a microplate reader (Spark, TECAN). Three independent experiments were conducted.

### Primary culture of neonatal mouse cardiomyocytes

C57 mice (SLAC Laboratory Animal Co. Ltd, Shanghai, China) were kept under standard animal room conditions (temperature 21±1 °C; humidity 55–60%) with food and water continuously available. Cardiomyocytes were isolated from 1-3 days old neonatal mouse hearts. Briefly, after hearts were digested by trypsin and Collagenase II, cells were suspended in DMEM with 15% FBS, and pre-cultured in humidified incubator (5% CO_2_) for 90 min to obtain cardiac fibroblasts for their selective adhesion. Then the suspended cardiomyocytes were plated in another dish. The purity of cardiomyocytes was increased by supplementing BrdU (5-Bromo-2’-deoxyuridine) to prevent non-cardiomyocytes from developing. Culture medium was renewed after 48 hours.

### Antibodies and Chemicals

Primary antibodies used to detect proteins were: BMP4 (ab235114, Abcam), KAT2A (ER63516, Huabio, Hangzhou, China), Histone H3 (acetyl K9) (HA722132, Huabio, Hangzhou, China), GAPDH (10494-1-AP, Proteintech), and β-actin (HA722023, Huabio, Hangzhou, China). secondary antibody: goat anti-rabbit IgG-h□+□HRP Conjugated (HA1023 Huabio, Hangzhou, China). Zinc sulfate heptahydrate was purchased from Sigma-Aldrich (Z0251). Vorinostat, Entinostat, Romidepsin, Pyrithione, and Zinpyr-1 were purchased from Med Chem Express (HY-10221, HY-12163, HY-15149, HY-B1747, and HY-D0155). ROS was detected using a Reactive Oxygen Species Assay Kit (CA1410, Solarbio, Beijing, China)

### RNA Isolation and Quantitative Real-time PCR (RT-PCR)

Total RNA was isolated from HL-1 cells, AC16 or neonatal mouse cardiomyocytes by using a TRIzol reagent (Takara Biotechnology, Dalian, China) according to the manufacturer’s instructions. 5 µg total RNA in a volume of 20 µL was reversely transcribed by using Hifair® ?1st Strand cDNA Synthesis SuperMix (11137ES60, Yeasen, Shanghai, China). After the termination of cDNA synthesis, mRNA levels of target genes were determined by qRT-PCR. Amplification was performed by CFX Opus 96 (Bio-Rad, USA). Real-time PCR was cycled in 95 °C/10 s, 60 °C/20 s and 72 °C/30 s for 40 cycles, after an initial denaturation step at 95 °C for 5 min using Hieff® qPCR SYBR Green Master Mix (11201ES08, Yeasen, Shanghai, China).The relative amounts of the mRNA levels of the target genes were normalized to the β-actin levels, respectively, and the relative difference in mRNA levels was calculated by 2−ΔΔCt method. The primers for quantitative Real-time PCR (RT-PCR) were listed in the Supplement Data (Table S1).

## Results

### Zinc induces the downregulation of *Bmp4* transcription in mouse myocardial cells

We conducted CCK-8 assays on mouse and human cardiomyocyte cell lines, HL-1 and AC16, respectively, using varying concentrations of zinc sulfate (ZnSO_4_) (Figure1A). To ensure maximal stimulation of the cells without disrupting their normal growth and metabolism, we chosen 70 μM as the experimental concentration. Next, 13 important genes related to cardiac development and growth-*Bmp4, Gata4, Hand1, Hand2, Mef2c, Notch1, Tbx1, Tbx5, Nkx2-5, Isl1, Wnt3a, Wnt2b*, and *Foxa2*-were selected as targets for transcriptional analysis [10, 11]. The results showed that under ZnSO_4_ treatment, the transcription of *Bmp4* and *Isl1* in HL-1 were significantly downregulated (Figure 1B). However, in AC16 cells, no significant changes were observed in the expression of these 13 genes (data not provided).

**Fig. 1.**
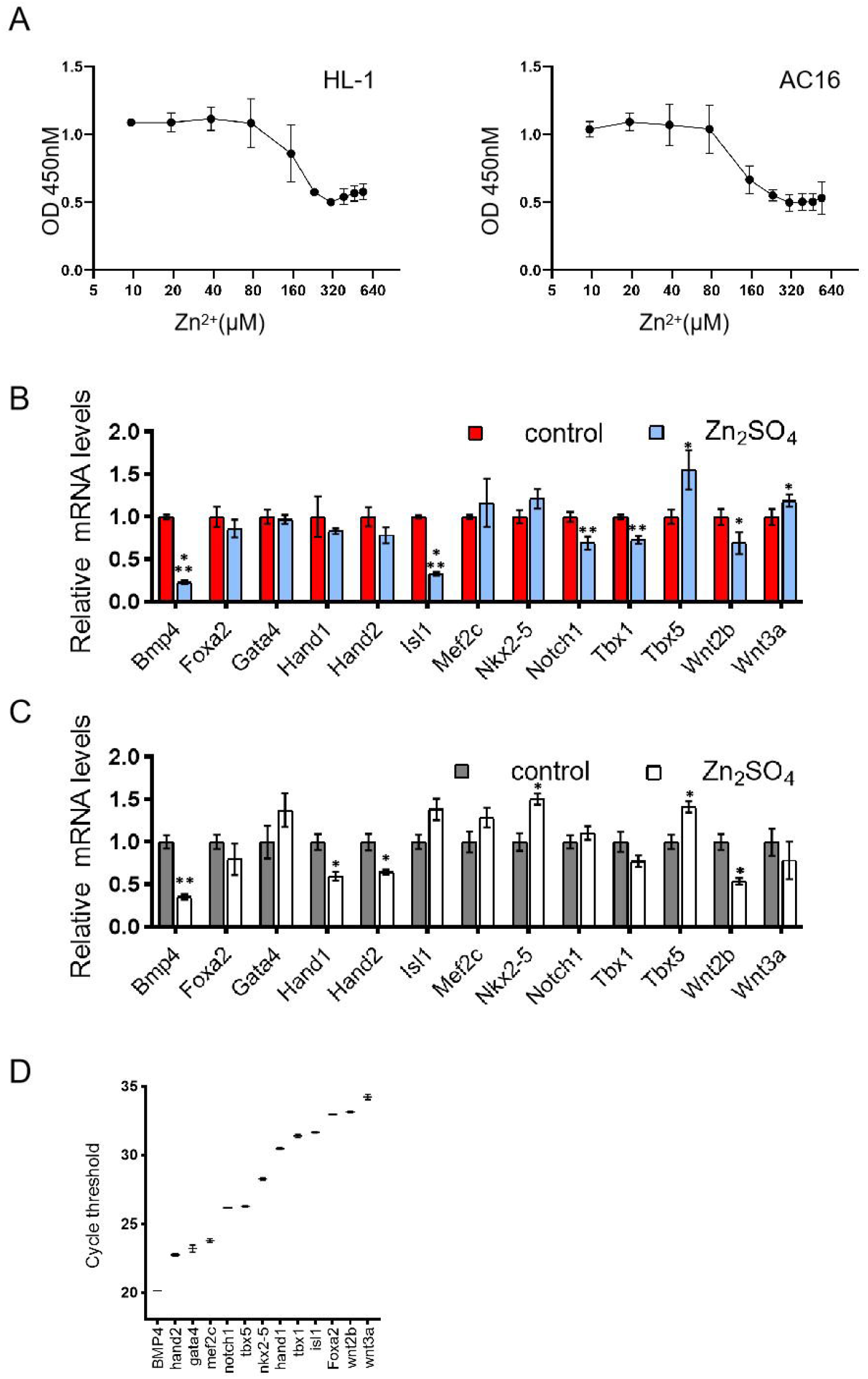
Zinc inhibits the transcription of *Bmp4* in mouse cardiomyocytes. (A) CCK-8 proliferation assay in HL-1 and AC16 cardiomyocytes. N=4. (B) HL-1 cells and (C) mouse primary cardiomyocytes were treated with 70 μM ZnSO_4_ for 24 hours. mRNA levels of 13 genes were analyzed. N=3. (D) The qPCR cycle thresholds of 13 gene in mouse primary cardiomyocytes with no treatment. N=3. All data are presented as means ± SD. *p < 0.05; **p < 0.01; ***p < 0.001 versus relevant control

Next, we collected primary cardiomyocytes from neonatal mice aged 1-3 days and cultured them in vitro. After treating primary cardiomyocytes with ZnSO_4_ for 24 hours, the transcriptional of *Bmp4* showed a significant downregulation, while other genes also exhibited certain degrees of changes (Figure 1C). Taking into account of the differential mRNA expression levels of 13 genes in neonatal mouse cardiomyocytes (Figure 1D) and the changes observed in HL-1 cells (Figure 1C), we selected *Bmp4* for further in-depth investigation.

We halved the concentration of ZnSO_4_ in the current experiment, and the inhibition of *Bmp4* mRNA expression was significantly alleviated, although a reduction was still observed (Figure 2A). This indicates that the inhibitory effect exhibits a notable concentration dependency. Also, the inhibitory effect was reversible. In the comparative sampling across multiple time points, we observed that *Bmp4* mRNA level significantly decreased as early as 4 h. The inhibition reached its maximum at 6 h and showed a gradual decline by 24 h.

**Fig. 2.**
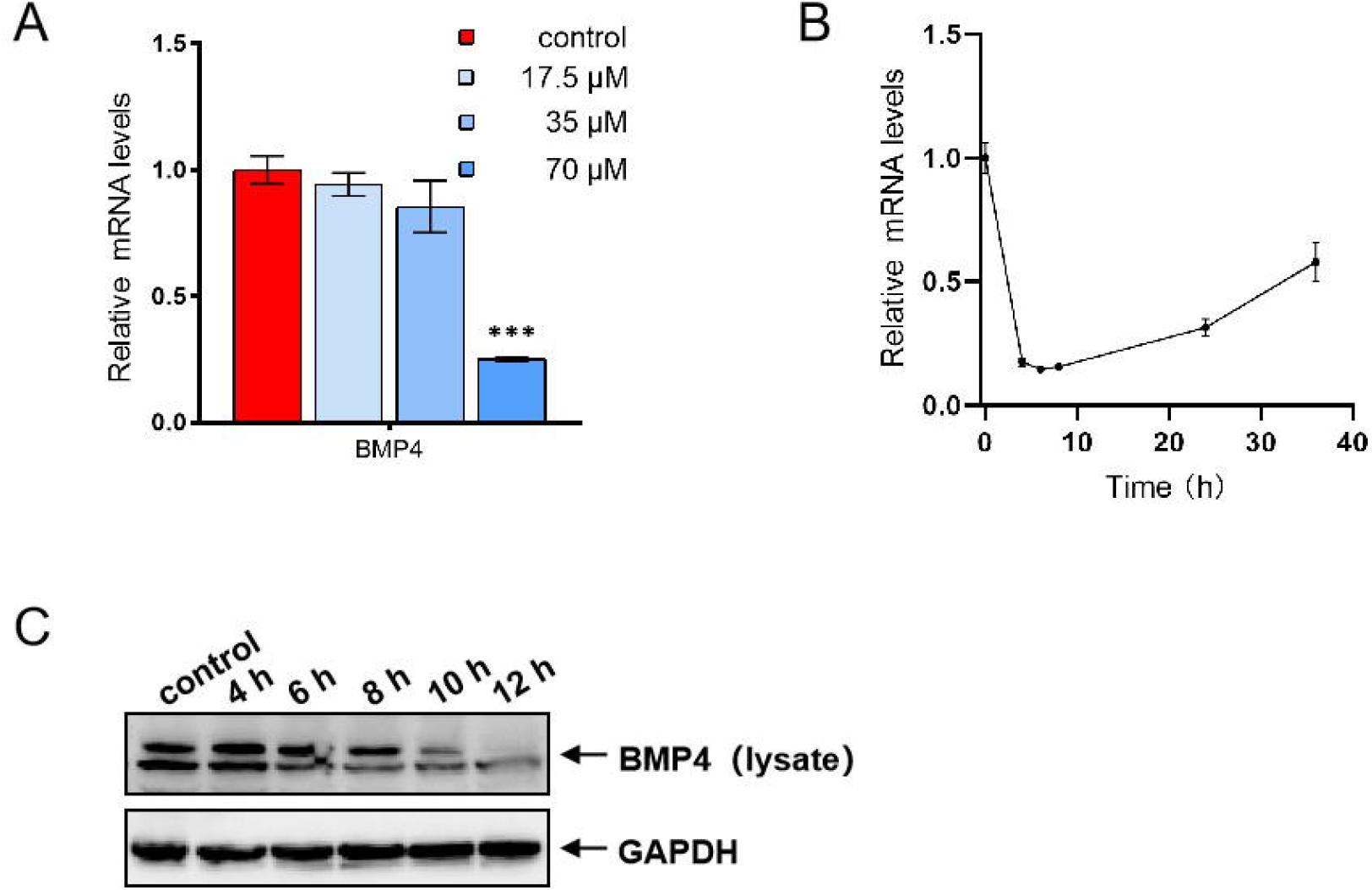
The inhibition of *Bmp4* in HL-1 cardiomyocytes by zinc is both time- and concentration-dependent. (A) mRNA levels of *Bmp4* under different concentrations of ZnSO_4_. N=3. (B) the transcriptional level changes of *Bmp4* over a period of 36 hours in the presence of 70 μM ZnSO_4_. (C) Immunoblotting analyses of BMP4 at different time points after treatment with ZnSO_4_. All data are presented as means ± SD. ***p < 0.001 versus relevant control

With an extended treatment duration of 36 hours, *Bmp4* mRNA level progressively recovered to more than half of its original (Figure 2B). In immunoblotting analysis, the inhibition of BMP4 by zinc lagged behind the transcriptional change, with discernible differences only becoming apparent at 6 h (Figure 2C).

### Zinc suppresses *Bmp4* expression through the reduction of histone acetylation

*Bmp4* is known to promote cardiomyocyte differentiation and is considered as an important cardiomyocyte marker [12]. Some studies suggest that both cardiac growth and maintenance of homeostasis are associated with covalent modifications of histone deacetylases (HDACs) [13-16]. Therefore, we utilized a broad-spectrum HDAC inhibitor, Vorinostat, to treat HL-1 in combination with zinc. The result indicates that Vorinostat is able to counteract the effects of zinc, thereby maintaining *Bmp4* mRNA at its original level in HL-1 cells (Figure 3A).

**Fig. 3.**
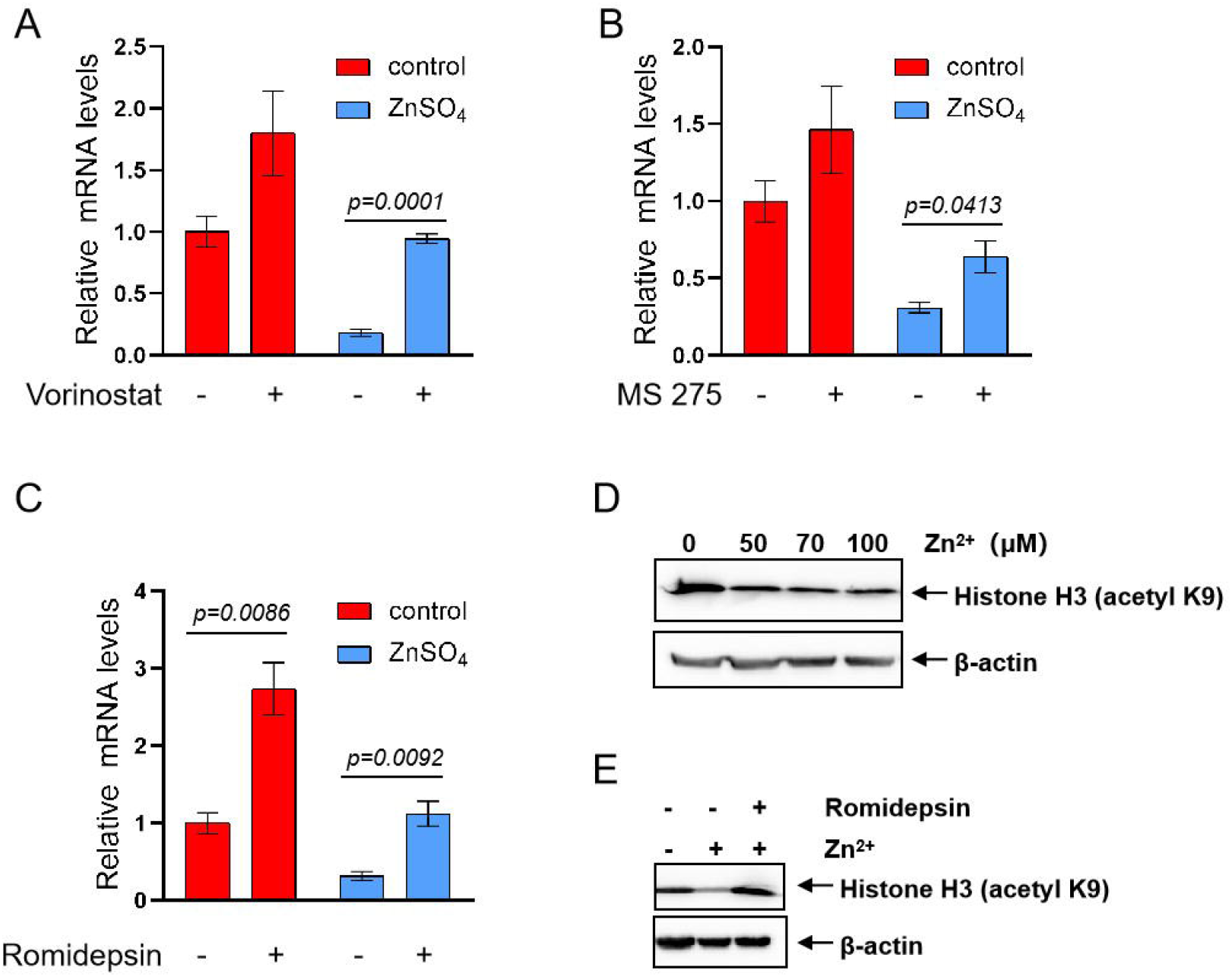
Zinc inhibits the transcription of *Bmp4* through reducing the acetylation of histones. (A) Vorinostat, (B) Entinostat, and (C) Romidepsin were all capable of reversing the downregulation of *Bmp4* induced by ZnSO_4_, with their working concentrations being 5 μM, 4 μM, and 50 nM, respectively. (D) Zinc significantly reduced H3K9ac. (E) Romidepsin significantly reversed H3K9ac decreased by zinc

This suggests that the transcription of *Bmp4* is regulated by histone acetylation. HDACs exert their catalytic function in conjunction with other proteins as part of a complex. And HDAC complexes exhibit specificity in their transcriptional regulation [17]. This infer that *Bmp4* is regulated by specific HDACs. The type I histone deacetylase (HDAC1,2,3) inhibitor Entinostat preliminarily confirmed this (Figure 3B). Next, we used a more targeted inhibitor, Romidepsin (HDAC1,2), and found that it significantly inhibited the downregulation of *Bmp4*. Moreover, when Romidepsin acted alone, it markedly increased the expression of *Bmp4* (Figure 3C).

Thus far, we have narrowed down the range of HDACs capable of regulating *Bmp4* to HDAC1/2. Both of them share a common catalytic site in histones at H3K9 [17] . Then the acetylation levels at H3K9 were tested. ZnSO_4_ greatly reduced H3K9ac even at 50 μM (Figure 3D). Furthermore, Romidepsin reversed the deacetylation effect at H3K9 caused by zinc (Figure 3E).

### Zinc reduces histone acetylation levels by downregulating the transcription of acetyltransferases

Histone acetyltransferases (HATs) and HDACs exert opposing effects and work together to maintain dynamic equilibrium in histone acetylation. After treating HL-1 with ZnSO_4_, we found no significant change in mRNA levels for 11 HDACs (Figure 4A). However, for acetyltransferases (*Kat2a, Kat2b, Kat6a, Kat7*, and *Kat14*) capable of producing H3K9ac,their transcription were all downregulated (Figure 4B). Immunoblotting confirmed that KAT2A expression was suppressed by zinc treatment (Figure 4C). Thus, we proposed that zinc disrupts the previously maintained balance in histone acetylation levels through downregulating HATs.

**Fig. 4.**
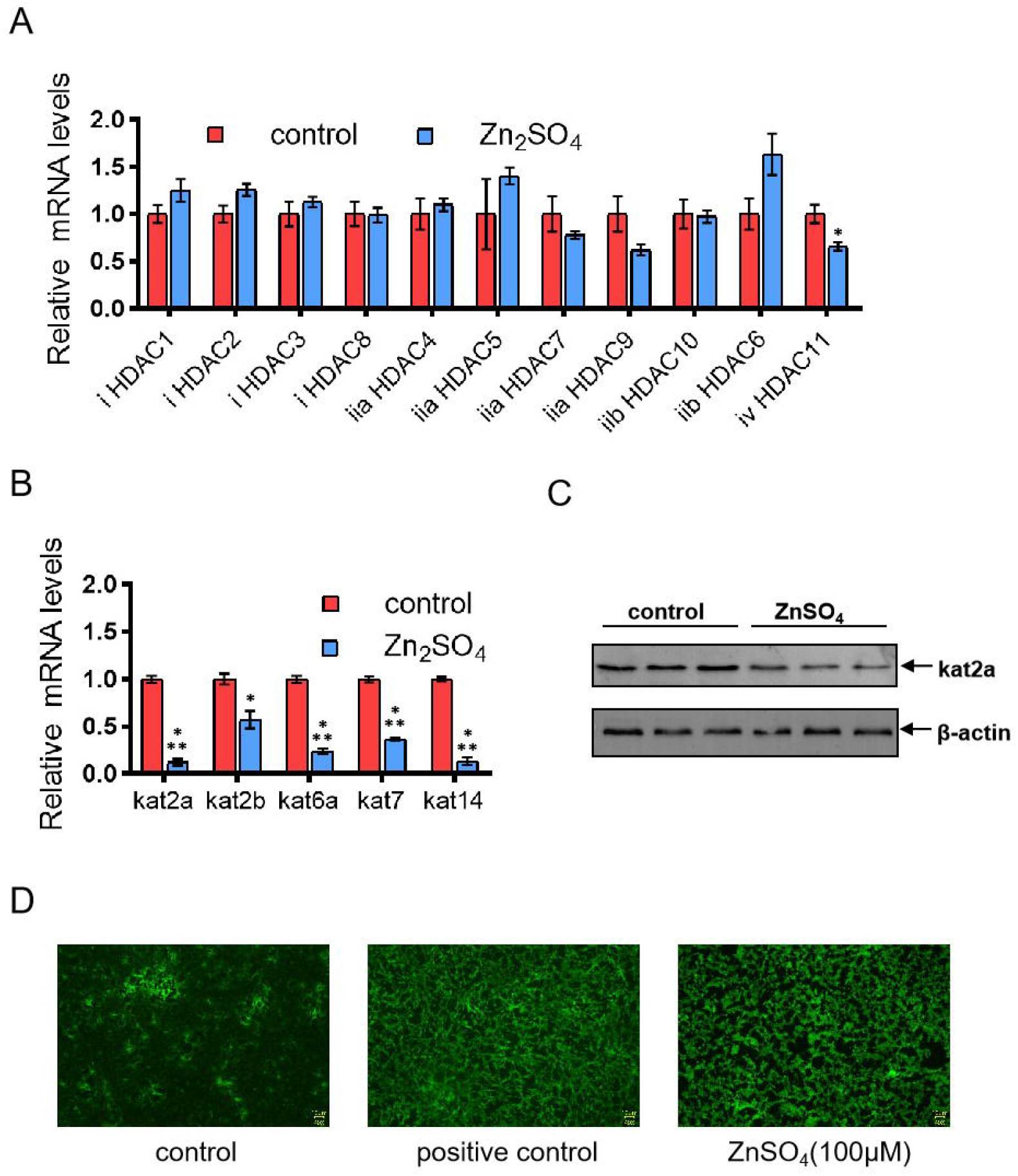
Zinc reduces histone acetylation by inhibiting the expression of HATs rather than HDACs. mRNA expressions of (A) 11 HDACs and (B) 5 HATs under treatment of ZnSO_4_. N=3. (C) Immunoblotting analyses of KAT2A under treatment of ZnSO_4_. (D) Zinc induced the generation of ROS in HL-1 cells. ZnSO_4_ (100 μM). Positive control, H_2_O_2_ (50 mg/L). All data are presented as means ± SD. *p < 0.05; ***p < 0.001 versus relevant control

Zinc has been reported to impede mitochondrial function, including excessive ROS, decreased pyruvate, and reduced ATP [5, 18]. As a downstream product of pyruvate, acetyl-CoA is the substrate for histone acetylation. Therefore, we suspected that the impairment of mitochondrial function may lead to a decrease in acetylation levels. Some researchers have proposed that zinc overload induced a substantial amount of ROS in rat cardiomyocyte H9C2 [19]. Similarly, zinc triggers ROS in HL-1 cells (Figure 4D). However, Hydrogen peroxide, which can induce mitochondrial oxidative stress and generate a large amount of ROS, was unable to downregulate *Bmp4*.

### Zinc indirectly inhibits histone acetylation

Zinc has been proposed to directly inhibit HDAC1 activity [20]. Moreover, zinc also has a certain degree of activating effect on acetyltransferases [21]. It should be noted that both studies were carried out in kit tests investigating the effects of zinc on enzyme activity from nuclear extract. In a different way, we simulated direct interaction of zinc with HATs and HDACs while preserving cellular integrity. To enable zinc ions to directly interact with acetylases and deacetylases in nucleus, we utilized the zinc ionophore pyrithione to facilitate the entry of Zn^2+^ into the cells without relying on zinc ion transporters. After the addition of pyrithione, Zn^2+^ at concentrations around the normal physiological range (10.71∼18.36 μM) caused severe cytotoxicity. Through optimization, we identified a relatively mild zinc concentration 1.5 μM for effective treatment. Nevertheless, the transcription of *Bmp4* was not downregulated; instead, it showed an increase. Notably, when the Zn^2+^ concentration was raised to 7.5 μM, *Bmp4* was upregulated more than 10 folds (Figure 5A). It was evident that under these conditions, intracellular free zinc ([Zn^2+^]i) has permeated throughout the entire cell (Figure 5B). Concurrently, the cell state has significantly deteriorated. For cells cultured in normal medium, [Zn^2+^]i remains predominantly localized in the cytoplasm, and a clear boundary between the cytoplasm and the nucleus is still maintained (Figure 5B).

**Fig. 5.**
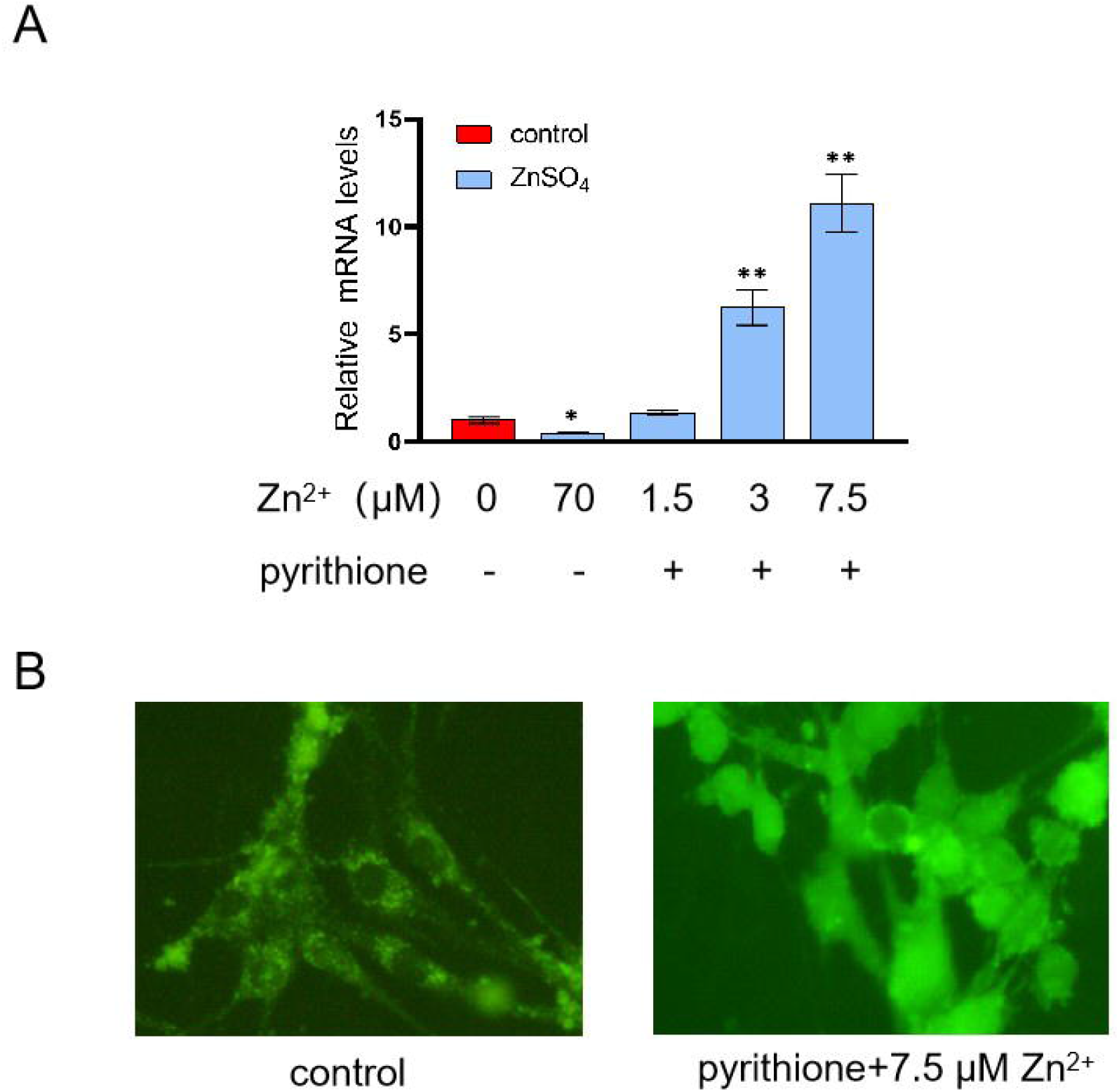
Zinc indirectly inhibits *Bmp4* transcription. (A) Zinc promotes the transcription of *Bmp4* with the assistance of pyrithione. (B) The distribution of free zinc ions in HL-1with or without pyrithione. Pyrithione, zinc ionophores, 10 µM. HL-1 cells labeled with 5 µM Zinpyr-1 for 0.5 h at 37 °C. All data are presented as means ± SD. *p < 0.05; **p < 0.01 versus relevant control

## Discussion

Under zinc treatment, the gene transcription changes in HL-1 and AC16 cells, both being cardiomyocytes, are not consistent. We believe this could be attributed to the limited scope of the genes studied. Thus, this does not necessarily mean that excessive zinc has no effect on human cardiomyocytes. Further, this discrepancy may be due to differences in genetic backgrounds, leading to variations in the chromosomal environment of the same gene, thereby resulting in differences in regulatory patterns. In an earlier study, *Bmp4* was demonstrated to be regulated by HDAC1/2 in mouse embryonic lung cells [22]. However, this regulatory mechanism becomes ineffective in the lung cells of postnatal mouse. Therefore, even for the same gene within the same cell, the regulatory mechanism may undergo changes over different periods.

The transcription of *Bmp4* is regulated by both HATs and HDACs. In our study, zinc downregulates the expression of HATs but does not alter the levels of HDACs. This combined effect results in a reduction of histone acetylation. Although the mitochondrial oxidative stress induced by hydrogen peroxide did not result in the downregulation of *Bmp4* in this study, changes in mitochondria, as organelles that provide acetyl groups for histone acetylation, remain noteworthy. The acetyltransferases have a broad impact on cellular processes.

But little is known about how these enzymes themselves are regulated. This study provides a model for research in this field.

Multiple zinc transporters work in coordination to maintain cellular zinc homeostasis. Under normal physiological conditions, the influx and efflux of zinc are tightly regulated by the Zrt- and Irt-like proteins (ZIP) family and the zinc transporter (ZnT) family, respectively [1]. During 36 hours of ZnSO_4_ treatment, the transcriptional expression of *Bmp4* in HL-1 cells initially decreased and then gradually recovered. This phenomenon may be attributed to the coordinated action of zinc transporters, which reduced intracellular zinc levels, thereby allowing histone acetylation levels to gradually recover. However, the addition of zinc ionophores disrupted this balance, reducing the barrier effect of cellular membrane on zinc ions. Consequently, the concentration of [Zn^2+^]i significantly increased. Zinc with high concentration directly inhibit the activity of the deacetylase HDAC1 [20]. Thus, the entire histone acetylation increased. Although the protein levels of acetyltransferases reduced at this stage, the histone acetylation level was still beyond the control. As a result, *Bmp4* transcription level was significantly elevated. To sum up, under normal physiological condition, excess zinc in the culture medium downregulates histone acetylation through an indirect way.

The concentration of zinc in extracellular body fluids (10.71–18.36 μM) differs significantly from that in the cytoplasm (pM–nM) [1, 23]. ZIPs and ZnTs maintain zinc homeostasis. Once their functions are dysregulated and zinc homeostasis is disrupted, many genes regulated by histone acetylation, including oncogenes and tumor suppressor genes[24], will exhibit diametrically opposite regulatory directions, Which will severely disrupt the normal physiological state of the cell. Therefore, the functional relationship between histone acetylation and zinc transporters warrants deeper attention and further research.

## Acknowledgments

This research was supported by the National Nature Science Foundation of China (NO. 82270309).

## Author Contribution

Shu Xu, Chao Tang, and Weize Xu designed the experiments. Shu Xu conducted the experiments and wrote the manuscript. Yuzhuang Hu provided discussions for the experiment.

## Data Availability

No datasets were generated or analyzed during the current study.

## Declarations

### Ethics approval

Animal experiments were approved (approval no. 21045) by the Animal Ethics Committee of Zhejiang University (Zhejiang, China) and performed according to the Guide for the Care and Use of Laboratory Animals (NIH Publication no. 85⍰23, revised 1996).

